# Spatially non-continuous relationships between biological invasion and fragmentation of mangrove forests

**DOI:** 10.1101/2021.04.16.440248

**Authors:** Zhen Zhang, Jing Li, Yi Li, Wenwen Liu, Yuxin Chen, Yihui Zhang, Yangfan Li

## Abstract

Rapid and large-scale biological invasion results in widespread biodiversity loss and degradation of essential ecosystem services, especially in mangrove forests. Recent evidence suggests that the establishment and dispersal of invasive species may exacerbated in fragmented landscape, but the influence of mangrove fragmentation on coastal biological invasion at landscape scale remains largely unknown. Here, using the derived 10-m resolution coastal wetland map in southeast coast of China, we examine the relationships between fragmentation of mangrove forests and salt marsh invasion magnitude and quantify the geographical variations of the relationships across a climatic gradient. Our results show that mangrove forests with small size, large edge proportion, and regular boundary shape tend to suffer more serious salt marsh invasions, indicating a positive correlation between mangrove fragmentation and its invaded magnitude. In particular, such fragmentation-invasion relationships in subtropics are shown to be more intensive than in tropic. Our findings provide the first spatially explicit evidence of the relationships between mangrove fragmentation and biological invasion on a landscape scale, and highlight an urgent need for conservation and management actions to improve mangrove connectivity, which will increase resistance to invasions, especially for small-size subtropical mangrove forests.

## 1. Introduction

In exotic species-introduce hotspots such as the coastal area, biological invasion is becoming a major driver of biodiversity decline and losses of ecological services in coastal forests (Bellard et al., 2016; Slingsby et al., 2017). As the dominated woody wetland community along tropical and subtropical coastlines, mangrove forests are particularly sensitive to invasive species due to the narrow habitat niche within the intertidal environment for species competition and intensive anthropogenic disturbances which limits their ability to migrate landward (Bradley et al., 2012). While only covering a small portion of the Earth’s surface, mangrove forests provide a diversity of essential ecosystem services far beyond the land area they occupy, such as coastal protection, carbon sequestration, erosion prevention, and habitat for fisheries (Donato et al., 2011; Worthington et al., 2020). However, the supply of these key ecosystem services has been threatening as mangroves are invaded by more than 50 plant species globally (Biswas et al., 2018). For example, woody mangroves invaded by herbaceous salt marshes reduce their carbon storage and slow down rates of sediment accretion in response to sea level rise (Kelleway et al., 2017). In addition to biological invasion, expansion in aquaculture, logging and coastal reclamation has led to unprecedented fragmentation of mangrove forests in recent few decades (Bryan-Brown et al., 2020; Richards & Friess, 2016). Fragmentation of mangrove communities may raise exposure to exotic species along forest edges (Dawson et al., 2015), further increasing their ecological sensitivity to invasive species. In the light of such potential intersection, efforts to understand the impacts of mangrove fragmentation on biological invasion have become critical for effective management action.

Growing evidence suggests that landscape fragmentation would promote the establishment of invasive species, and resident native species in fragment landscape are more likely to expose to invader (Malavasi et al., 2014; Vilà & Ibáñez, 2011). For example, positive correlations have been reported between landscape fragmentation and exotic species richness, as well as abundance in both tropical rainforest and subtropical dry forest (Aguirre-Acosta et al., 2014; Waddell et al., 2020). This fragmentation-invasion relationship can partly be interpreted as transformation of biophysical environments near habitat fragment edges (Ordway & Asner, 2020). However, this was concluded in terrestrial ecosystems, illustrating our poor knowledge on the relationships between coastal mangrove fragmentation and biological invasions. In contrast to terrestrial forests, mangroves could be invaded by both aquatic and terrestrial species (Biswas et al., 2018). Additionally, limited habitats in intertidal environments and relatively smaller size of mangrove patches imply that fragmentation in mangrove patches may have more complicated consequences contrast with terrestrial forests. As such, conclusion derived in terrestrial forests could not be directly transformed to coastal mangrove forests. A better understanding of fragmentation-invasion relationships in mangrove ecosystems is required to warn potential invaded mangrove communities, and also to perform efficient management strategies in the context of anthropogenic land-use change.

The establishment and spread of alien species are affected by both biotic factors and abiotic conditions (Liu et al., 2018; van Kleunen et al., 2018), including species richness of native communities (Beaury et al., 2020), competitive and consumptive interactions (Alofs & Jackson, 2014), resident herbivores (Zhang et al., 2018) and local temperature (Cornelissen et al., 2019). Due to the spatial heterogeneity of these conditions, invasive ability of alien species and resistant of native species to invasion may not spatially continuous across the landscape (Stotz et al., 2016). As a result, invasion processes and outcomes may vary among different climate zones such as tropic and subtropics. Understanding how this geographical variation affect invasion pattern thus is fundamental to identify vulnerable areas where are particularly sensitive to invasion, as well as developing conservation strategies in advance. However, earlier studies on fragmentation-invasion relationships mostly conduced in site level, can be sensitive to the local environmental conditions (Oehri et al., 2017), constrained by their poor spatial coverage while ignoring the geographical variations of fragmentation-invasion relationships. While some studies monitored the spatial distribution of invasive species in a larger scale by using remote sensing approaches (Liu et al., 2018; Vaz et al., 2018), there is still uncertainties about how the fragmentation-invasion relationship varies across a climatic gradient.

Here, we adopt a biogeographic perspective to test whether the mangrove fragmentation has impacts on the invasion magnitude of an invasive saltmarsh species (*Spartina alterniflora*) along latitudinal gradient in Southeastern China, a region that has undergone widespread *S. alterniflora* invasion over the last several decades (Meng et al., 2020). We characterize mangrove fragmentation from three key aspects, including fragment size, edge, and shape, which have been widely used in fragmentation researches (Haddad et al., 2015; Mendes et al., 2016). The main purposes of this study are: (a) to test the latitudinal pattern in invasion magnitude of *S. alterniflora* on mangrove forests, (b) to specify the impacts of mangrove fragmentation on *S. alterniflora* invasion magnitude, and (c) to detect the geographic variations in fragmentation-invasion relationship along climatic gradient.

## 2. Materials and Methods

### 2.1. Study Area

The Southeast coastal areas of China provide an ideal setting to examine the invaded pattern of native wetland plants on a landscape scale as the area has suffered from a long-term *S. alterniflora* invasion and is one of the most sensitive mangrove-marsh ecotones in the world to changes in climatic conditions (Osland et al., 2016). *S. alterniflora* was introduced and spread over a 19°-latitude region along the east coast of China since 1979 (Liu et al., 2016), and dominated a large area of tidal flats in past two decades, occupying approximately 545.80 km^2^ of coastal zone in 2015 (Liu et al., 2018). This study was conducted within the coastal areas in the southeast of mainland China, which spans a latitudinal gradient extending from tropical Leizhou Peninsula in the south (∼20°34′N) to subtropical Zhejiang Province (∼28°25′N), covering the whole geographical range of mangrove-salt marsh ecotone in mainland China (Chen et al., 2017).

### 2.2. Characterizing mangrove fragmentation

#### 2.2.1. Fragment size

We mapped the distribution of mangroves and *S. alterniflora* in 2020 using all available atmospherically corrected Sentinel-2 surface reflectance product on Google Earth Engine platform (Gorelick et al., 2017). The derived coastal wetland dataset at 10-m resolution enables a high-accuracy assessment of relationship between mangroves and *S. alterniflora* (88.7% overall accuracy; see Supplementary Information for more details). Since the fragmentation process is inevitably resulted from the reduction of the patch area, we used the mangrove fragment size as a metric to measure fragmentation (Haddad et al., 2015). The size (unit: ha) of each mangrove fragment was then calculated based on the derived wetland dataset using an Albers conic equal-area projection.

#### 2.2.2. Edge proportion

A mangrove fragment is composed of edge and core area (Fig. 1). Mangrove edge is defined as any mangrove pixel adjacent to a non-mangrove pixel, and the mangrove core is the remaining mangrove pixels (Chaplin-Kramer et al., 2015). The specific process of edge area identification was carried out using a sum filter with a three by three window (8 neighbors) on the binary mangrove/non-mangrove image, in which mangrove pixels were assigned with value of 1 and the non-mangrove pixels with value of 0. Mangrove pixel was considered as a grid of ‘core area’ only if its value equals to 9 based on the spatial filter process; otherwise, it was classified as ‘edge area’. We calculated the edge proportion in each mangrove fragment (i.e. the ratio of edge area to mangrove fragment area), and used it as a fragmentation metric to explore the edge effect of mangrove fragments to *S. alterniflora* invasion.

**Fig. 1.**
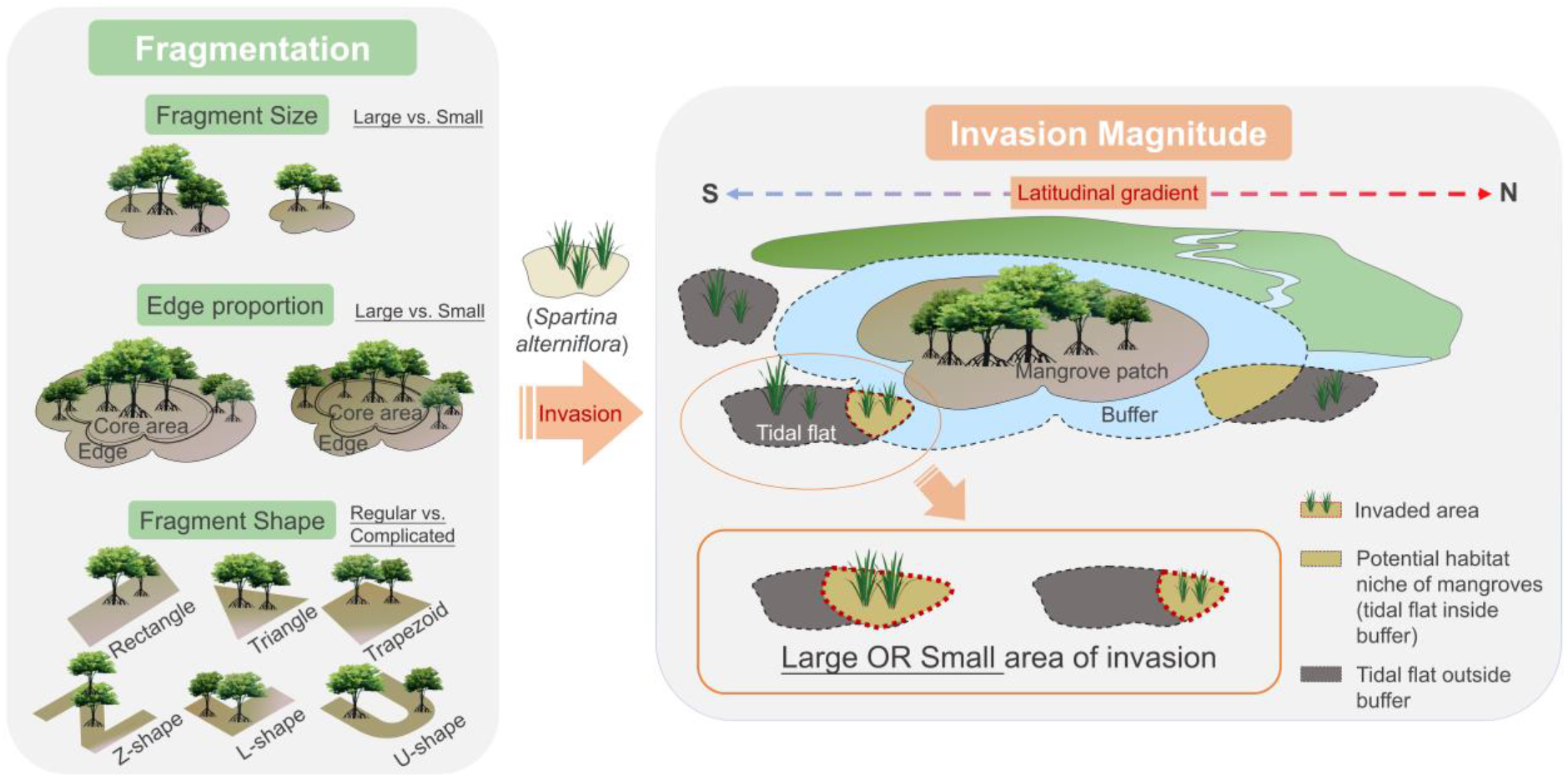
Conceptual diagram of seascape patterns of mangrove fragments and the *S. alterniflora* invasion magnitude.

#### 2.2.3. Fragment shape

We proposed an information theory-based approach to identify the specific shape of grid-cell mangrove fragments (Supplemental Fig. S2; see Supplementary Information for more details). To group all fragments into certain shapes, we predefined six standard shapes as rectangle, triangle, trapezoid, Z-shape, L-shape, and U-shape, since these shapes can approximately describe most of our pixel-based mangrove fragments according to visual inspection. The first three shapes were classified as regular shapes and the others as complicated shapes for extracting conclusive information (Supplemental Fig. S3). The reason we use specific shapes rather than shape complexity to reflect the shape effect is that specific shapes are practical in guiding mangrove restoration.

For each mangrove fragment, the distances of centroid to fragment boundary measured from all directions were plotted as a continuous signal (Supplemental Fig. S2). Jensen-Shannon (JS) divergence (Lin, 1991), an information theory-based metric to assess the similarity between two signals, was then used to compare the signal of each mangrove fragment with signal of standard shapes. By comparing the identified signal of each fragment and that of standard shape, we assigned every mangrove fragment to certain shape group, which can be further grouped to regular and complicated shapes. All spatial related processes were conducted in Python 2.7, and more details about our information theory-based approach are provided in Supplementary Information.

### 2.3. Quantifying *S. alterniflora* invasion magnitude

*S. alterniflora* tend to occur on the bare tidal flats close to the margins of mangrove forest or within the canopy gaps of sparse mangroves (Chen et al., 2016). Our earlier experiment (Zhang et al., 2012) demonstrated *S. alterniflora* transplanted into the understory of dense mangrove stands died in a period of time, implying that the invasion of *S. alterniflora* did not directly occupy areas of mangroves, but the surrounding bare tidal flats of mangrove forests to limit the mangrove propagation. Thus, the *S. alterniflora* invasion magnitude was defined as the proportion of surrounding tidal flats occupied by *S. alterniflora* to the total bare tidal flats around mangrove forests (Fig. 1).

Under this circumstance, we firstly determined which bare tidal flats have the potential to support mangrove propagation based on the historic expansion distance of mangrove forests. Specifically, we calculated the historical expansion distance of mangroves during the past two decades using the multi-temporal global mangrove maps from the Global Mangrove Watch dataset (Bunting et al., 2018). Considering the capability of mangrove propagation would be spatially heterogeneous, we calculated the distance per 1° latitudinal band, and used it as the radius to create buffer area for related mangrove fragments. Bare tidal flats in created buffer zone were identified as potential habitat for mangrove propagation. Subsequently, based on the global tidal flat dataset (Murray et al., 2019) and our *S. alterniflora* map, we recorded the tidal flats area and *S. alterniflora* area within the surrounding potential habitat separately, and calculated the ratio of the *S. alterniflora* area to the area of tidal flats as the measure of invasion magnitude. The calculated invasion magnitude was bounded between value 0 and 100%, and a higher number means that more of the tidal flats where mangroves have the potential to propagate has been invaded by *S. alterniflora*.

### 2.4. Statistical analysis

#### 2.4.1. Detecting latitudinal patterns

To analyze the detailed latitudinal variations of invaded magnitude and fragmentation of mangrove communities, we averaged these metrics in each of the 0.1° latitudinal bands. We then used ordinary least squares regression models to estimate the biogeographical trend in invasion magnitude and fragmentation metrics along latitudinal gradient, and the significance of the models were assessed. Considering the shape index of mangrove fragments is a qualitative variable, and all mangrove fragments were divided into two groups (regular and complicated shapes), we calculated the proportion of mangrove fragments in both groups per 0.1° latitudinal band, and used it to represent latitudinal variation in fragment shapes. Additionally, to examine the difference of invasion magnitude in different climate zones, we compared *S. alterniflora* invasion magnitude between the tropical and subtropical mangrove fragments by using dependent *t*-test at fragment scale.

#### 2.4.2. Effects of fragment size on invasion magnitude

To specify the effect of mangrove fragment size on *S. alterniflora* invasion, we aggregated mangrove fragments with similar sizes into same group by using *K*-means algorithm and obtained 83 and 60 fragment groups for tropical and subtropical regions, respectively. Aggregating mangrove fragments of similar size into groups, regardless of location and other factors, allowed us to isolate the effect of fragment size on biological invasion (Hansen et al., 2020). Due to the observed nonlinear relationship, we conducted a piecewise regression (Toms & Lesperance, 2003) and asymptotic regression (Stevens, 1951) to model the nonlinear size effect respectively, and identified two change points derived from these regression models. We assessed the support for the change points by comparing the Akaike information criterion (AIC) of the piecewise model and asymptotic model with the AIC of simple linear model. The mean values of change points were defined as the thresholds to divide mangrove fragments into “small” (below threshold) and “large” (above threshold) (Ordway & Asner, 2020). Because the response variable (i.e., average *S. alterniflora* invasion magnitude) ranged between value of 0 and 100%, beta regressions were then applied on both small and large size of mangrove fragments to model the relationship between mangrove size and *S. alterniflora* invasion proportion. For examining the impacts of macroclimatic condition on fragmentation-invasion relationships, we conducted the above statistical analysis for tropical and subtropical mangroves separately.

#### 2.4.3. Edge effect on invasion magnitude

Mangrove fragments in similar edge proportion were grouped through *K*-means analysis, and a group-level statistical analysis was applied to isolate the effect of fragment edge on biological invasion. We conducted a beta regression model on average invasion magnitude and edge proportion to investigate the edge effect on invasion. Considering the edge proportion may be affected by fragment area, we added the average fragment size into beta regression model as an explanatory variable to account for this interaction effect.

To estimate the geographic variation of edge effect on *S. alterniflora* invasion, we further investigated the latitudinal variation of edge effect. For each 0.1°latitudinal band, we regressed invasion magnitude and edge proportion on fragment scale, and defined the edge sensitivity of mangroves to invasion as the slope of the regression models. Preliminary graphical exploration of the relationship between latitude and statistically significant edge sensitivity was conducted to provide details on likely landscape-level patterns. Observed tipping latitude was then identified using piecewise regression model. To investigate potential mechanisms behind the latitudinal variation of edge effect, we used the Normalized Difference Vegetation Index (NDVI) as a proxy of mangrove canopy cover and aboveground biomass (Myneni et al., 1997) to explore the latitudinal trend in mangrove structures.

#### 2.4.4. Shape effects on S. alterniflora invasion

Analysis of variance (ANOVA) and Tukey honestly significant difference (HSD) post-hoc test were conducted to explore the impacts of mangrove fragment shape on *S. alterniflora* invasion. Considering the bias of different sizes of mangrove fragment in same shape, we divided all mangrove fragments into three categories by size through *K*-means algorithm: small-size (< 0.7 ha), medium-size (0.7-24.4 ha) and large-size (> 24.4 ha). Afterwards, we applied a univariate ANOVA on each category to estimate differences among all mangrove shapes, and an HSD post-hoc test to identify the types of shape that related to the least invasion of *S. alterniflora*. Similar analysis was conducted on mangrove fragments which were divided into three groups according to edge area and core area. All statistical analyses were implemented in R version 4.0.0 (R Development Core Team, 2020) using the packages ‘changepoint’ (Killick & Eckley, 2014), ‘segmented’ (Muggeo, 2008) and ‘betareg’ (Cribari-Neto & Zeileis, 2010).

## 3. Results

### 3.1. Latitudinal patterns of *S. alterniflora* invasion magnitude

According to the distribution of mangroves and *S. alterniflora* derived from remote sensing images, there was 172.46 km^2^ of mangroves and 181.27 km^2^ of *S. alterniflora* in study area (Fig. 2). The variability of the area of mangroves and *S. alterniflora* was observed across the latitudinal gradient (Fig. 2b). As the latitude rises, area of mangrove decreased while that of *S. alterniflora* increased. A significant shift of the mangrove-*S. alterniflora* dominance along latitude were observed near the Tropic of Cancer (23°26′N). In the tropical regions (south of Tropic of Cancer), the total area of mangroves (158.65 km^2^) was nearly eight times of *S. alterniflora* (21.83 km^2^); while in the subtropical regions, the dominance plant was replaced by *S. alterniflora* (159.44 km^2^), which was nearly 12 times of mangroves about 13.81 km^2^.

**Fig. 2.**
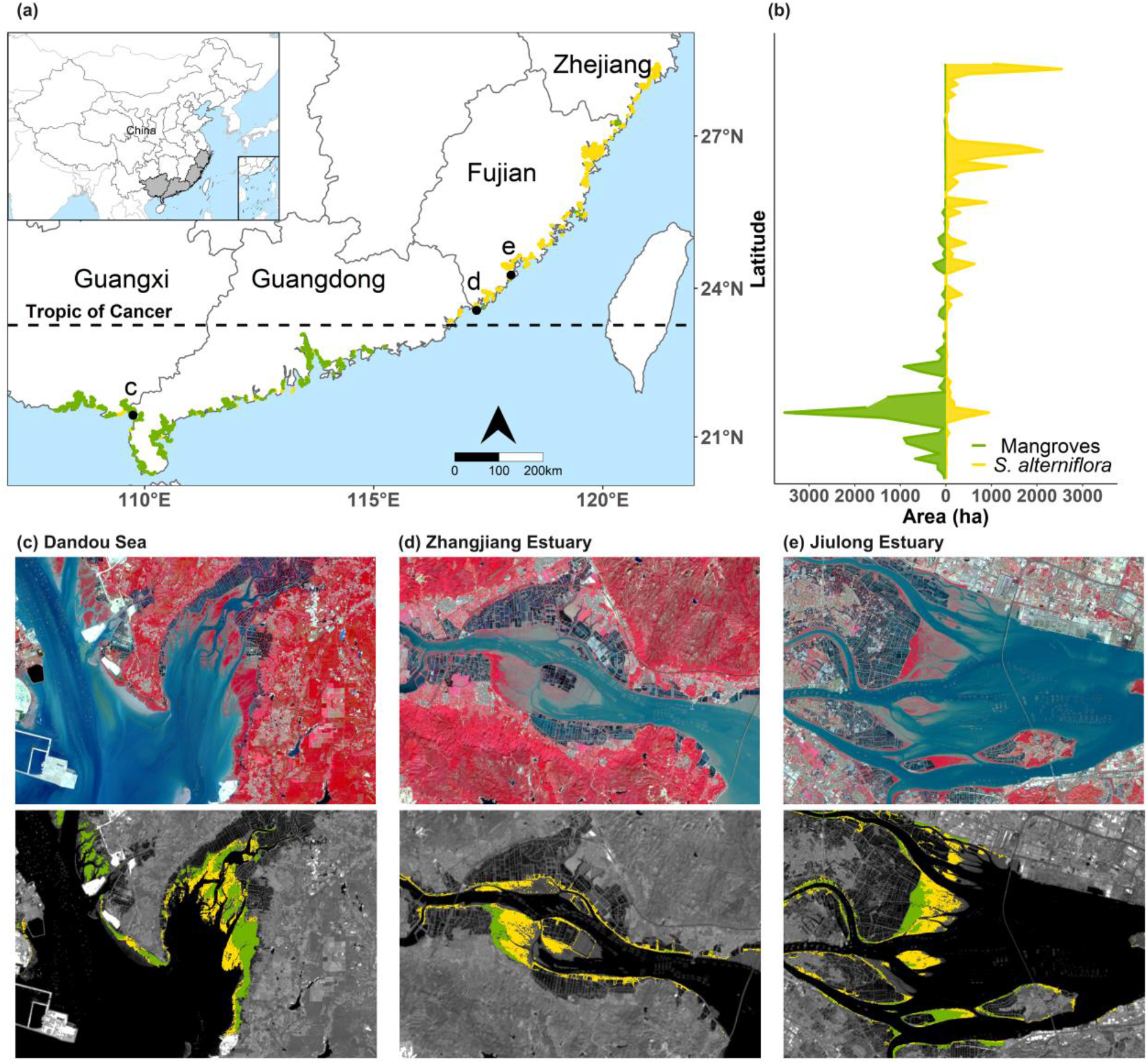
Spatial distribution of mangroves and *Spartina alterniflora* in 2020. Spatial distribution with 0.1°latitude summaries of mangrove and *S. alterniflora* area are shown as (a) and (b). Three examples of spatial patterns of mangroves and *S. alterniflora* were shown as (c) Dandou Sea, (d) Zhangjiang Estuary, and (e) Jiulong Estuary.

Major invasion of *S. alterniflora* to mangroves was observed across the study region with a significant increasing trend along latitude from 20°34′N to 28°25′N (Fig. 2 and 3a). Subtropical mangroves had a mean value of 34.0% invaded area by *S. alterniflora*, and the tropical mangroves was about 29.2% (*t*-test, *p* < .001; Fig. 3b). The distribution of *S. alterniflora* invasion suggests that mangroves in subtropical region were more vulnerable in terms of invasion than their tropical counterparts, which was consistent with the spatial distribution of mangrove-*S. alterniflora* dominance.

**Fig. 3.**
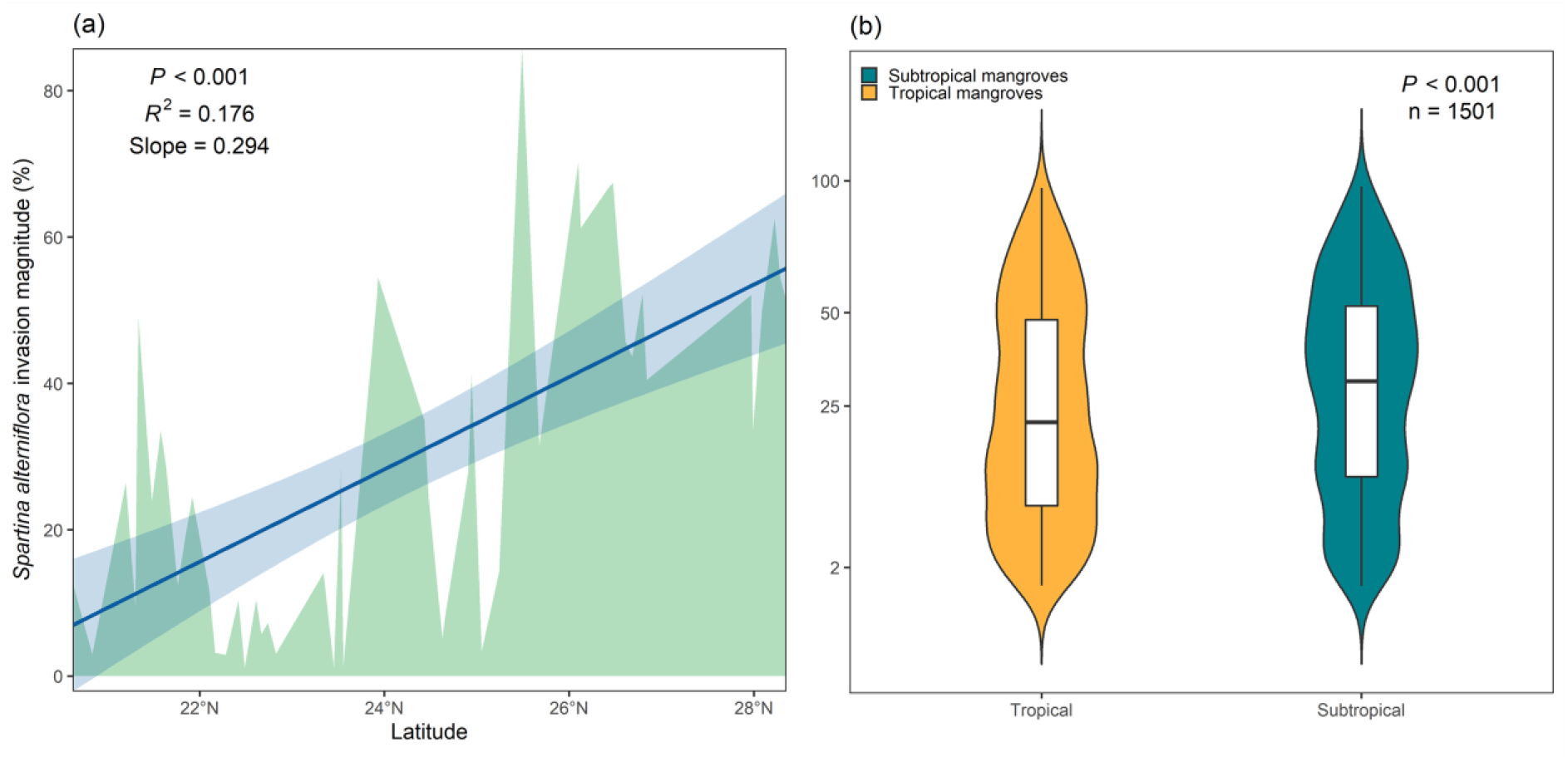
Latitudinal variation in the invasion magnitude of *Spartina alterniflora* showing as (a) latitudinal profile and (b) comparison between subtropics and tropic. The observed total amount of invaded mangrove fragments is 1501 (751 and 750 in subtropical and tropical regions, respectively). Blue line in (a) represents linear latitudinal trend of invasion magnitude, and blue shaded area is the 95% confidence interval. Green shaded area represents the distribution of invasion magnitude along latitudinal gradient. The boxes in (b) show data within the 25th and 75th percentile, black lines show the median values of each group, and violin-shaped area represents distribution of invasion magnitude values. The *y* axe in (b) was square root-transformed as necessary to comply with parametric assumptions, but show untransformed values.

### 3.2. Latitudinal variations in mangrove fragmentation and its effects on invasion

#### 3.2.1. Fragment size

The distribution of average mangrove fragment size follows a decline trend across the latitudinal gradient (Supplemental Fig. S4a), in consistence with the latitudinal trend of total mangrove area. A one-degree increases in latitude resulted in an approximately 0.25 ha decrease of average mangrove fragment size (*R*^2^ = .101, *p* = .010). Both piecewise regression and nonlinear asymptotic regression indicated potential nonlinearity in the relationships between mangrove fragment size and invaded magnitude (supported over linear regression based on lower AIC, ΔAIC = 2.5 and 3.7 for piecewise model and asymptotic model in tropic, and ΔAIC = 52.1 and 55.3 for piecewise model and asymptotic model in subtropics). By combining these two nonlinear models, we found thresholds of mangrove size at the value of 3.93 ± 1.12 and 4.44 ± 0.61 ha for tropical and subtropical mangroves, respectively (Fig. 4). Size of mangrove under 4.44 ha in subtropical region was inversely correlated with *S. alterniflora* invasion (Pseudo *R*^2^ = .483, *p* < .001); however, there is no relationship between the *S. alterniflora* invasion and mangrove size once the mangrove fragment is larger than the threshold value of 4.44 ha (Pseudo *R*^2^ = .041, *p* = .206). For tropical mangroves, the relationship between fragment size and *S. alterniflora* invasion magnitude was not statistically significant both in small (Pseudo *R*^2^ = .002, *p* = .705) and large mangrove fragments (Pseudo *R*^2^ = .001, *p* = .834).

**Fig. 4.**
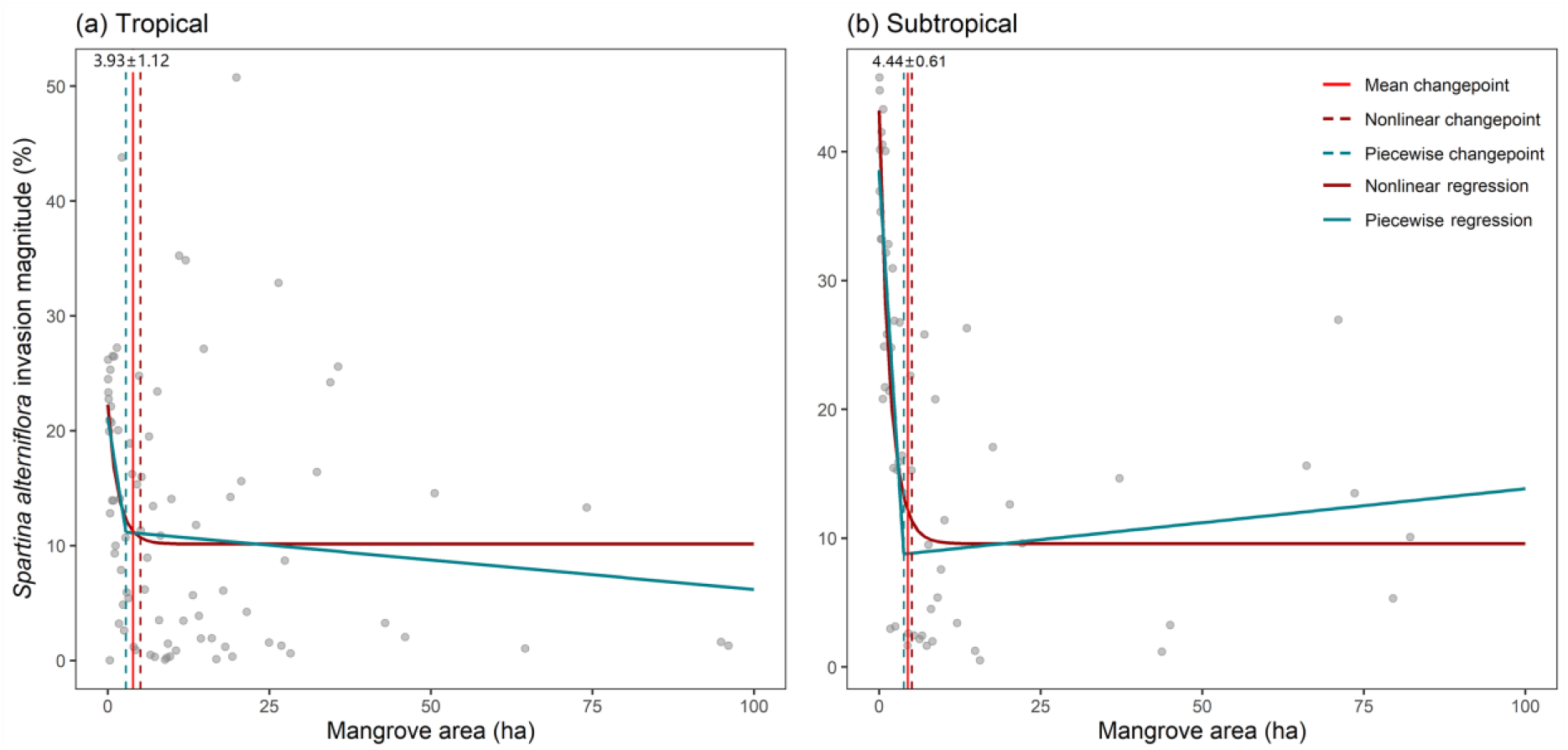
Effects of mangrove fragment size on invasion magnitude of *Spartina alterniflora* in tropical (a) and subtropical (b) areas. Values between dashed lines are the detected interval of abrupt changes in invasion when size of mangrove fragment increased in certain values. Detailed model parameters are shown in Supplemental Table S3.

#### 3.2.2. Edge proportion

The average edge proportion in each group classified by mangrove edge proportion was related linearly to invasion magnitude of *S. alterniflora* (Fig. 5a), suggesting that mangrove fragments with large proportion of edge belt were particularly sensitive to *S. alterniflora* invasion (Pseudo *R*^2^ = .561, *p* < .001). Mangrove fragments with more than 75% of edge area (979 out of 1501 fragments) experienced an average invasion of 34.2%; 263 mangrove fragments with edge proportion of 50-75% showed an average *S. alterniflora* invasion of 26.2%; mangrove forests with 25-50% edge proportion (174 fragments) showed an invasion magnitude of 16.5%; and only 85 mangrove forests with the smallest edge proportion (< 25% of edge proportion) were exposed to the lowest invasion magnitude of 10.1%. The positive effect of mangrove edge proportion on *S. alterniflora* invasion magnitude was modified by an interaction between edge proportion and size (ΔAIC = 1.2 comparing the full linear model with or without interaction), with edge effect will be minimized when the mangrove fragment size increases (slope = -0.001, *p* = .036, Supplemental Table S4).

**Fig. 5.**
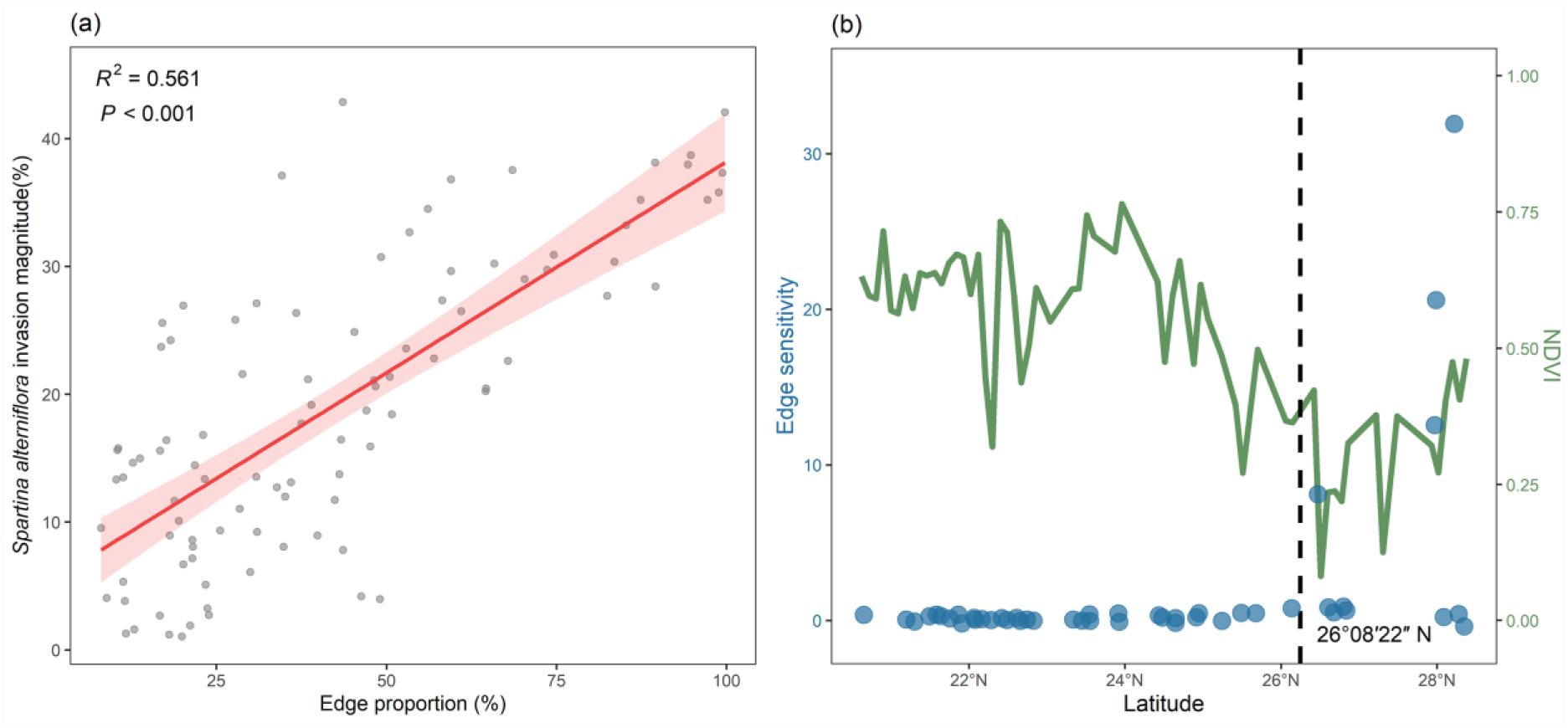
Edge effects of invasion magnitude. (a) Relationship between edge proportion and *Spartina alterniflora* invasion magnitude. The shaded area indicates the 95% confidence interval of regression. (b) Edge sensitivity (regression slope of edge effect) and NDVI of mangroves along latitude (average value of mangrove forests in each 0.1°latitudinal band). The vertical dashed line shows the latitudinal change point (26°08′22″ N).

As the edge proportion of mangrove fragments increases with latitude, the increased edge effect implies that subtropical mangroves are particularly sensitive to *S. alterniflora* invasion (Supplemental Fig. S4b). Observing the edge sensitivity of mangroves across latitude through a piecewise model (ΔAIC = 5.2 comparing with the linear model), we detected a latitudinal change point at the location of 26°08′N (95% Confidence Interval (CI) = 25°16′ to 27°22′; Fig 5b), adjacent to the northern limit of natural mangrove distribution in China (27°20′N). The latitudinal fluctuation of mangrove NDVI provides a direct estimation of changes in mangrove canopy cover and biomass (Fig. 5b), which explained the phenomenon of abrupt shift of edge effect on invasion across the latitudinal tipping point. It exhibited a latitudinal heterogeneity in the north part of this change point, suggesting that the resistance of planted mangroves to *S. alterniflora* invasion was more susceptible to change of edges.

#### 3.2.3. Fragment shape

Different to former two metrics, latitudinal trend of mangrove fragment shape was not significant (Supplemental Fig. S4c). Comparing the *S. alterniflora* invasion in different shapes of mangroves, we found that invasion magnitude varied significantly in different-shaped fragments (*p* < .05; Fig. 6a). Regarding the small (< 0.7 ha) and medium size (0.7-24.4 ha) of mangroves, fragments with regular shapes of triangle, trapezoid and rectangle appeared to have an average invasion magnitude of 31.78% ± 1.82% (95% CI), which was substantially larger than mangrove fragment with complicated U, L and Z-shapes by 26.20% ± 1.81% (95% CI). But the effect of different shapes in large-sized mangroves (> 24.4 ha) was not observed (*p* = .681; Supplemental Table S5).

**Fig. 6.**
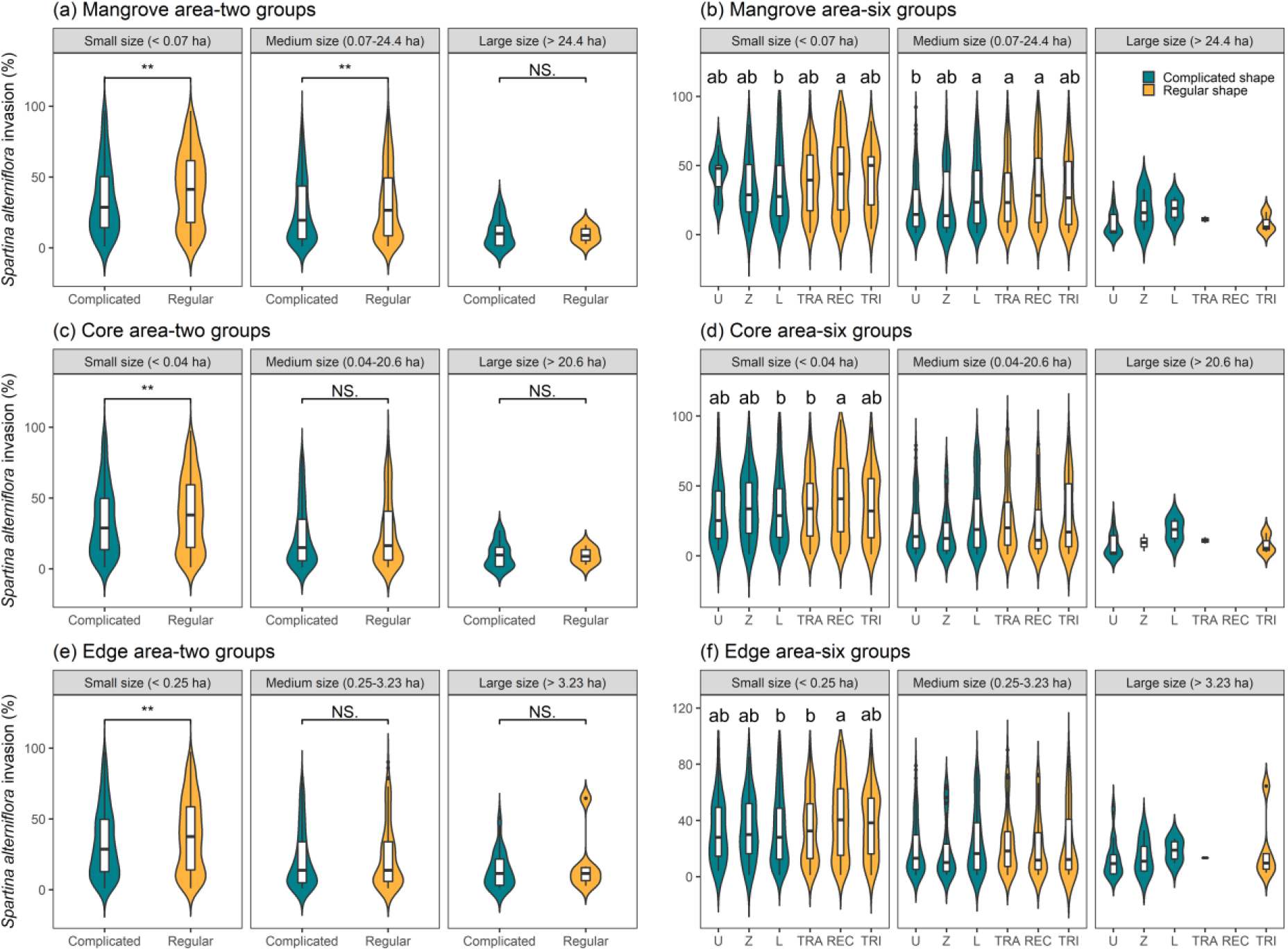
Comparing *Spartina alterniflora* invasion by size (a, b), core area (c, d) and edge area (e, f) of mangrove fragments. Values of *S. alterniflora* invasion area were indicated by violin plot regarding different shapes of mangrove fragment. Welch’s *t*-test was used to estimate differences between complicated and regular-shaped mangroves. One-way ANOVA was conducted to evaluate the differences of invasion among all six shapes. Boxes in the middle of each dataset show the interquartile ranges, and solid lines in the box indicate the median value of such dataset. Letters above violin plot bars denote significant difference among shapes resulting from multiple comparison analysis. Different letters indicate significant differences (*p* < .05) among mangrove fragment shapes. Individual statistical values for each subplot are presented in Supplemental Table S5. Series with double asterisk (**) means the correlation analysis of *p* < .01, and NS. stands for non-significant at the level of *p* > .05. Abbreviation of shapes are list as TRA-trapezoid, REC-rectangle, TRI-triangle.

In addition, we conducted similar analysis on different-shaped mangroves with considering different edge and core area of each mangrove fragment (Fig. 6c, e). In terms of invasive magnitude, we observed a significant difference between regular and complicated shapes only in small-size groups with edge area < 0.25 ha and core area < 0.04 ha. This finding indicates the impact of mangrove fragment shapes on *S. alterniflora* invasion existed in the case of fragmented mangrove communities with small-sized core area and edge area.

The largest invasive magnitude was observed in the rectangle-shaped (regular shape) mangrove fragments, which was higher than trapezoid-shaped (regular shape) mangroves by 7.33% ± 6.87% (95% CI) and L-shaped (complicated shape) mangrove fragments by 9.80% ± 8.21% (95% CI; Fig. 6b; Supplemental Table S5). The small size of L-shaped mangrove fragments had the most stable structure to minimize *S. alterniflora* invasion, which signifies the L-shaped mangrove fragments have a high level of resistance in invasion challenges of *S. alterniflora*.

## 4. Discussion

Spatial detection of the mangrove-*S. alterniflora* ecotone is of great value to understand dynamics of biological invasion in coastal wetlands. This study, to our knowledge, is the first to examine relationships between mangrove fragmentation and its invaded magnitude at a landscape scale. We found that fragmentation of mangrove forests would promote the invasive magnitude of *S. alterniflora*, and this relationship varies between latitudes and climate zones. Our findings highlight the value of large scale and biogeographic perspective on mangrove fragmentation and biological invasion, with the goal of informing policy and ecosystem management in the future.

### 4.1. Latitudinal trends in mangroves and *S. alterniflora*

Our study indicates that the invasion magnitude of *S. alterniflora* on native mangroves increased with increasing latitude, suggesting the mangrove community may be more susceptible to invasion at higher than at lower latitudes. This response is consistent with biogeographical patterns of their climatic niche, and agrees with previous study which indicated the aboveground biomass of *S. alterniflora* increases with latitude in Southern China (Liu et al., 2020). In low latitudes, climate conditions are suitable for survival and dispersal of freeze-sensitive mangroves, enhancing their competition on habitat niche with invasive salt marsh (Cavanaugh et al., 2019). Conversely, the expansion of mangrove forests in subtropical and temperate regions is limited by its climate conditions, which in turn facilitate the invasion of freeze-tolerant salt marshes (Osland et al., 2017). Another possible explanation of this latitudinal pattern may be that the species richness of mangrove communities may decrease as the latitude rises, weakening the biotic resistance to invaders (Beaury et al., 2020; Guo et al., 2020; Wu et al., 2018).

### 4.2. Fragmentation-invasion relationships in mangrove-salt marsh ecotone

The negative nonlinearity in size effect implies that small mangrove fragments are particularly sensitive to *S. alterniflora* invasion as compared to large mangrove blocks. While the size effect is not significant in tropic, the constrain effect of size on relationship between edge proportion and invaded magnitude is significant on whole ecotone. Our findings reveal that large mangrove fragments exhibit stronger resistance to salt marsh invasion. This is consistent with previous studies indicating that large and highly connected patches could enhance dispersal ability and system’s robustness to stochastic perturbations (Wintle et al., 2019). Although the loss of small mangrove fragments may seem less concerning than large-scale invasion, their interconnectedness with adjacent habitats allows them to provide substantial ecosystem services as stepping stones to link local systems (Curnick et al., 2019). In addition, the invasion of small-sized mangroves would create barriers to species that depend on mangroves, erodes local coastal resilience and pushes mangrove ecosystems toward collapse. Therefore, protecting small mangrove fragments and preventing further mangrove fragments split into several smaller ones becomes the critical task in maintaining resilience in coastal ecosystems.

Our study highlights that the edge proportion of mangrove fragments was related with the incidence of plant invasions. Ecological effects arising from edges between mangrove and non-mangrove habitat change biophysical environments for adjacent habitats, generating the local conditions in which invasive species seem to thrive, such as canopy gaps and areas with high light levels (Li et al., 2014). Previous field experiment-based study also revealed that mangrove edge creates an open and low canopy structure in local community, which reduces herbivore activity but increases light availability, resulting in erosion of resistance of mangrove forests to *S. alterniflora* invasion (Zhang et al., 2018). We found that edge sensitivity of mangroves to invasion dramatically shifts at the location of 26°08’N (95% CI = 25°16’ to 27°22’N), which is near with the northern limit (27°20’N) of natural mangroves distribution (Chen et al., 2017), and is consistent with latitudinal variation of mangrove NDVI. This consistent geographic change point reveals that small fragmented mangroves with open canopy and planted mangroves are more susceptible to invasions.

Comparisons between multiple mangrove shapes suggests that the shape of mangrove fragments has significant impacts on salt marsh invasion, particularly in small and medium-sized mangrove fragments. Since there is no significant latitudinal trend in the shape of mangrove fragments, it is inferred that the shape effects on invasion may occur in most regions. Previous theory has noted that an ecologically optimum fragment shape tends to has a large core with some curvilinear boundaries and narrow lobes (Forman, 1995; Moser et al., 2002), but have yet to point specific shapes which have higher resistance to invasion. Here we report that mangrove fragments with regular shapes were more susceptible to *S. alterniflora* invasion, and rectangle was identified as the shape undergoing more severe salt marsh invasion and lower resistance. This can be explained by the resistance of mangrove fragments to external perturbations from nature environmental, is usually controlled by core area and interaction with adjacent habitats, both of which are determined by the shape of source fragments (Orrock et al., 2011). For testing the explanatory of core area on shape effect, we conducted an analysis of covariance (ANCOVA), which could control variable of total area in shapes that related with low invasion, and found that trapezoidal and L-shaped mangrove fragments have larger proportion of core area than that of rectangle (Supplemental Fig. S5). Additionally, mangrove fragments with trapezoid or L shape tend to have a large perimeter and size, thus these types of mangroves have a large biotic and abiotic flow with adjacent habitats and higher dispersal ability (Lester et al., 2007). As a result, fragment shapes with sufficient core area and curvilinear boundaries, such as L shape and trapezoid, exhibit stronger resistance to biological invasion (Rastandeh & Pedersen Zari, 2018).

### 4.3. Management implications for mangrove forests

Several global conservation policy mechanisms have included mangroves in their targets, such as the Mangroves for the Future and Sustainable Development Goals 14.5 and Target 11 in Aichi Targets (Friess et al., 2019). Regional governments have also taken actions into mangrove conservation. For example, Chinese government has established twenty-eight Protection Areas, which has protected 67% of the mangrove forests in China (Wang et al., 2020). The key way to achieve these goals is mangroves planting. However, results have shown large-scale replant planning did not lead to expected long-term mangrove area increases (Lee et al., 2019). In China, about half of the replanted mangroves have failed to restored, which is partly due to the invasion of *S. alterniflora* into suitable mangrove niche (Chen et al., 2009). Management and conservation of mangroves are therefore required for alleviating biological invasion stresses, which can be achieved by controlling landscape patterns of mangrove fragments in local communities according to our findings. The edge and shape sensitivity of mangroves to *S. alterniflora* invasion underpins effective guidance in rehabilitating and protecting existed mangrove fragments to improve their resistance to invasions. Moreover, preventing extant mangrove blocks divide into multiple smaller fragments should be a management priority wherever possible, especially for the subtropical mangroves due to their critical vulnerability to exotic species invasion. Paying more attentions to the size and edge proportion of replanted mangrove forests and prevent further mangrove fragmentation are crucial to mangrove conservation, especially in the regions beyond the geographic range limit of natural mangroves distribution.

Optimum fragment shape also needs to be considered in mangroves rehabilitation and management. The L- and trapezoid-shaped mangrove are recommended in this study as suitable fragment shapes to resist the invasion of alien species. Current mangrove restoration mainly focuses on natural environment, which neglects spatial structure and pattern of mangrove forests. Spatial characters, such as fragment size, edge proportion and fragment shape, should be incorporated to generate more effective restoration approach to enhance the resistance and resilience of mangroves and regional biosecurity.

### 4.4. Limitations

Although we presented observational evidences on the relationships between mangrove fragmentation and *S. alterniflora* invasion magnitude, there is still uncertainties associated with them as they were developed under some limitations. First, fragmentation metrics (fragment size, edge proportion and fragment shape) were calculated from raster dataset, which can be biased by its spatial resolution even if we use the highest resolution available remote sensing data (i.e., Sentinel-2 at 10-m resolution). Another limitation is except for the spatial distribution of the tidal flats, salinity gradient, soil texture, and tidal range may also affect the extent of potential mangrove habitat. These environmental factors can hardly be included in our analysis due to the lack of high-resolution data at the landscape scale.

## 5. Conclusions

By combining remote-sensing detection and landscape analysis, we demonstrate that fragmentation of mangrove forests increased the invaded magnitude by *S. alterniflora*, and this effect varies with latitudes and climate zones. We found that mangrove fragments with small size, large edge proportion and regular boundary shape are particularly sensitive to plant invasion. This has important conservation implications because mangroves are facing threats from both fragmentation and biological invasion globally. Our results indicate that the fragmentation-invasion relationships are ubiquitous throughout the whole mangrove-salt marsh ecotone, but intensified in subtropical regions, the inappropriate climatic area for freeze-sensitive mangroves. These findings suggest an urgent need for management strategy to mitigate fragmentation of mangrove forests, particularly in subtropical mangroves, and highlight that optimizing landscape pattern should be considered as an effective strategy for addressing the ongoing biological invasions, and can be used to navigate native species management actions for making management efforts more informed and effective.

## Declarations

## Acknowledgements

This research was supported by the National Natural Science Foundation of China (NSFC) Grant No.41701205 and Fundamental Research Funds for the Central Universities No.20720190089.

## Availability of data and material

All datasets used in this study are publicly available. The Sentinel-2 L2A Surface Reflectance (https://developers.google.com/earthengine/datasets/catalog/COPERNICUS_S2_SR); The Global Mangrove Watch dataset (https://data.unep-wcmc.org/datasets/45); the Global Tidal Flats dataset (https://www.intertidal.app/).

## Code availability

The code is available from https://github.com/GIS-ZhangZhen/Edge-sentivity-of-mangroves.

## CRediT authorship contribution statement

**ZZ**: Conceptualization; Methodology; Software; Formal analysis; Writing - Original Draft; Visualization

**JL**: Methodology; Formal analysis; Visualization

**Yi L**: Conceptualization; Writing - Review & Editing; Resources; Funding acquisition; Project administration

**WL**: Conceptualization; Formal analysis; Writing-Review & Editing

**YC**: Conceptualization; Formal analysis; Writing-Review & Editing

**YZ**: Conceptualization; Writing - Review & Editing

**Yangfan L**: Conceptualization; Resources

